# How light at night sets the circalunar clock in the marine midge *Clunio marinus*

**DOI:** 10.1101/2024.04.30.591642

**Authors:** Carolina M. Peralta, Eric Feunteun, Julien Guillaudeau, Dušica Briševac, Tobias S. Kaiser

**Author notes:** Shared senior authorship (Dušica Briševac, Tobias S. Kaiser). To whom all correspondence should be addressed: Dušica Briševac and Tobias S. Kaiser; Max Planck Institute for Evolutionary Biology, Max Planck Research Group Biological Clocks, August-Thienemann-Strasse 2, 24306 Plön, Germany.

## Abstract

Many organisms inhabiting the interface between land and sea have evolved biological clocks corresponding to the period of the semilunar (14.77d) or the lunar (29.53d) cycle. Since tidal amplitude is modulated across the lunar cycle, these circasemilunar or circalunar clocks allow organisms to adapt to the tides. Biological clocks are synchronized to external cycles via environmental cues called *zeitgebers*. Here, we explore how light at night sets the circalunar and circasemilunar clocks of *Clunio marinus*, a marine insect that relies on these clocks to control timing of emergence. We first characterized how moonlight intensity is modulated by the tides by measuring light intensity in the natural habitat of *C. marinus*. In laboratory experiments, we then explored how different moonlight treatments set the phase of the clocks of two *C. marinus* strains, one with a lunar rhythm and one with a semilunar rhythm. Light intensity alone does not affect the phase or strength of the lunar rhythm. Presenting moonlight during different 2-hour or 4-hour windows during the subjective night shows that (1) the required duration of moonlight is strain-specific, (2) there are strain-specific moonlight sensitivity windows and (3) timing of moonlight can shift the phase of the lunar rhythm to stay synchronized with the lowest low tides. Experiments simulating natural moonlight patterns with moonrise and moonset confirm that the phase is set by the timing of moonlight. Light intensity within the ranges observed in the field leads to the best synchronization. Taken together, we show that there is a complex and strain-specific integration of intensity, duration and timing of light at night to precisely entrain the lunar and semilunar rhythms. The observed fine-tuning of the rhythms under the most natural moonlight regimes lays the foundation for a better chronobiological and genetic dissection of the circa(semi)lunar clock in *C. marinus*.

## Introduction

The celestial movements of the earth and the moon entail regular changes in the environment, such as night and day, the seasons, the lunar cycle and the tidal cycle. Organisms have evolved corresponding biological clocks, i.e. endogenous time-keeping mechanisms, which are highly adaptive for each particular habitat, as they permit species to anticipate regular changes and time behavior or physiology accordingly. Moreover, they allow populations of individuals to synchronize important life history events such as reproduction. For each of the described geophysical cycles, there is a corresponding biological clock. The circadian clock (24 h) is the most widespread across the animal kingdom and by far the best-studied (Dunlap, 1999; Kaiser and Neumann, 2021). However, many organisms inhabiting the interface between land and sea have evolved biological clocks that allow them to adapt to the tides. These include the circatidal clock (12.4 h), the circasemilunar clock (14.77 d) and the circalunar clock (29.53 d) (reviewed in Neumann, 2014). Circasemilunar and circalunar clocks help organisms to adapt to the tides, because the tidal amplitude is modulated across the lunar cycle, being highest during the spring tide days just after full moon and new moon.

In this study we focus on the circasemilunar and circalunar clock of the marine insect *Clunio marinus*. *C. marinus* larvae and pupae settle in the lower intertidal, where they are almost constantly submerged (Neumann, 2014). However, the adults need the larval habitat to be exposed in order for oviposition to be successful. Exposure of the larval habitat occurs reliably every month during the spring tides, when tidal amplitude is highest. Therefore, *C. marinus* has a short adult lifespan of few hours, which is only used for reproduction and which is timed exactly to spring tide low tides. This is achieved via a combination of circasemilunar/circalunar and circadian clocks. The circalunar or circasemilunar clock synchronizes development and maturation with the spring tide days. Notably, there are *C. marinus* strains with a circalunar clock which only emerge during full moon or new moon, and there are strains with a circasemilunar clock which emerge during both full and new moon (Kaiser et al., 2021).The circadian clock gates adult emergence to one of the two daily low tides.

Biological clocks are synchronized to external environmental cycles via reliable timing cues, called *zeitgebers*. Once the zeitgeber effectively entrains (synchronizes) the clock, the endogenous rhythm will have a stable phase-relationship to the zeitgeber, the so-called phase of entrainment (Aschoff and Pohl, 1978). Thus, the zeitgeber is crucial for the synchronization of physiological processes to the external environmental cycles as it conveys information about the phase to the biological clock (Naylor, 2010).

The relevance of moonlight as an effective zeitgeber for lunar rhythms was first described in 1960 in the annelid *Platynereis* (Hauenschild, 1960). Since then, moonlight as a zeitgeber has been validated for many other species such as the brown algae (*Dictyota dichotoma* (Bünning and Müller, 1961), the crab *Sesarma haemotocheir* (Saigusa, 1980) and different *Clunio* species (*C. marinus*, *C. mediterraneus* and *C. tsushimensis*, reviewed in Neumann 2014). While in this study we focus on entrainment by moonlight, it is noteworthy that the zeitgebers for setting *Clunio*’s circalunar clock also include tidal cycles of mechanical agitation (Neumann, 1978) and tidal temperature cycles (Neumann and Heimbach, 1979; reviewed in Kaiser and Neumann, 2021).

Previous studies in *Clunio* started to explore how moonlight – or light at night – entrains the circalunar clock. It was shown that one night of continuous artificial moonlight is insufficient to properly synchronize a semilunar strain of *C. marinus* (Neumann, 1976). This led to the suggestion that light perception might be combined with some sort of counting mechanism (Neumann, 1987). Three, four or six nights with artificial moonlight are sufficient to entrain the circasemilunar or circalunar clocks (Neumann, 1966). Another major result was that a circadian clock regulates the perception of light at night (Neumann, 1989). *C. tsushimensis* was exposed to four consecutive days of constant darkness (DD) and artificial moonlight at different times of the subjective day and night. This has shown that only light given in the subjective night effectively entrains the circalunar clock (Neumann, 1995). The fact that this regulation of sensitivity also works under DD suggests that it is controlled by a circadian clock. Further studies on which particular times at night set the phase were performed by providing windows of light at night in shorter periods over the night. Exposing *C. tsushimensis* to a light dark-cycle of 12:12 and 3h of light at night shows that only moonlight in the middle of the night works as an effective zeitgeber (Neumann, 1987). Exposing the semilunar Santander (San) strain of *C. marinus* to 4h of light at night, suggests that moonlight is only perceived in the second half of the night (Neumann, 1969).

The phase relationship between artificial moonlight and emergence peak can be modified by changing different components of the artificial moonlight such as the number of successive nights with light at night (Neumann, 1976), varying the period of the light-dark cycle (Neumann, 1989; Neumann and Heimbach, 1979) and by a more complex simulated moonlight program (Neumann, 1985). The latter includes the daily shift in the rise and fall of the moon, as well as three different levels of light intensity corresponding to different lunar phases. When the tides were mimicked by superimposing an early, middle or late gate on the complex simulated moonlight program, the adult emergence peak did not follow the brightest moonlight, but shifted with the timing of light at night (Neumann, 1985). Finally, laboratory strains of *C. marinus* from different geographical locations show a strain-specific phase relationship between the artificial moonlight stimulus and the emergence peak in the laboratory (Neumann, 1966). It was shown that there is a good correlation between the time from the light stimulus to the emergence peaks in the laboratory and the time from midnight low-tides (when moonlight is presumably best perceived) to emergence in the field (Kaiser et al., 2011). Thus, *C. marinus* populations from different geographic origins show local adaptation in the phase relationship between moonlight perception and adult emergence.

Open questions remain, such as if and how light is modulated by the tides and to what extent the windows of light sensitivity vary across populations of *C. marinus*. It is also unclear which components of the moonlight – intensity, duration or daily timing – set the phase of emergence. In this study, we show by field measurements how moonlight intensity reaching the intertidal is modulated by the tides. In corresponding laboratory experiments we explored how different moonlight treatments set the phase of the circa(semi)lunar clock. We assessed two laboratory strains which have a lunar or a semilunar rhythm. We show that there is a complex and strain-specific interaction of intensity, duration and timing of light at night in setting the strength and phase of entrainment.

## Methods

### Light in the intertidal zone

In order to evaluate moonlight modulation by the tides in a *C. marinus* habitat, a RAMSES-ACC-VIS hyperspectral radiometer (TriOS GmbH) was placed in the lower littoral zone during the low tide of spring tides at Rocher de Bizeux in Dinard (see coordinates in TableS1, FigureS1). This site is near St. Briac-sur-Mer, from which a *C. marinus* strain has been obtained and described (Bria-1SL, Kaiser et al., 2021). The radiometer measured 192 wavelengths between 317-953 nm, at a resolution of ∼3 nm (data published on Max Planck Repository Edmond, https://doi.org/10.17617/3.6OXUES). Light was recorded every five minutes, over nearly four consecutive lunar months (106 nights) from 17-10-2013 to 01-02-2014. All timestamps were recorded in the time zone in which the device was deployed, in Central European Summer Time (CEST), and then converted to Coordinated Universal Time (UTC).

To investigate moonlight properties, we looked at changes in light intensity during the night over several months. As light intensity was low (i.e. for 502 nm on a full moon night, intensity was between 9.99×10^-^6-0.001 mW/m^−2^/nm^−1^), raw values were averaged in 30-minute intervals and values at each measured wavelength were summed in 100-nm bins. The highest light intensity detected in the middle of the night was of 0.06 mW/m^−2^/nm^−1^, on full moon (night between 17-11-2013 and 18-11-2013). Values above this threshold were considered as daylight and used to define the limits between night (Figure S2a-b) and day (Figure S2c-d).

The strongest intensity and clearest moonlight signal were observed between November 2013 and January 2014 (Figure S2a-b). Therefore, we subsetted the data for this time-frame (from new moon 03-11-2013 to new moon 02-01-2014, Figure 1a). In further text we referred to this dataset as “Dinard-underwater”.

**Figure 1.**
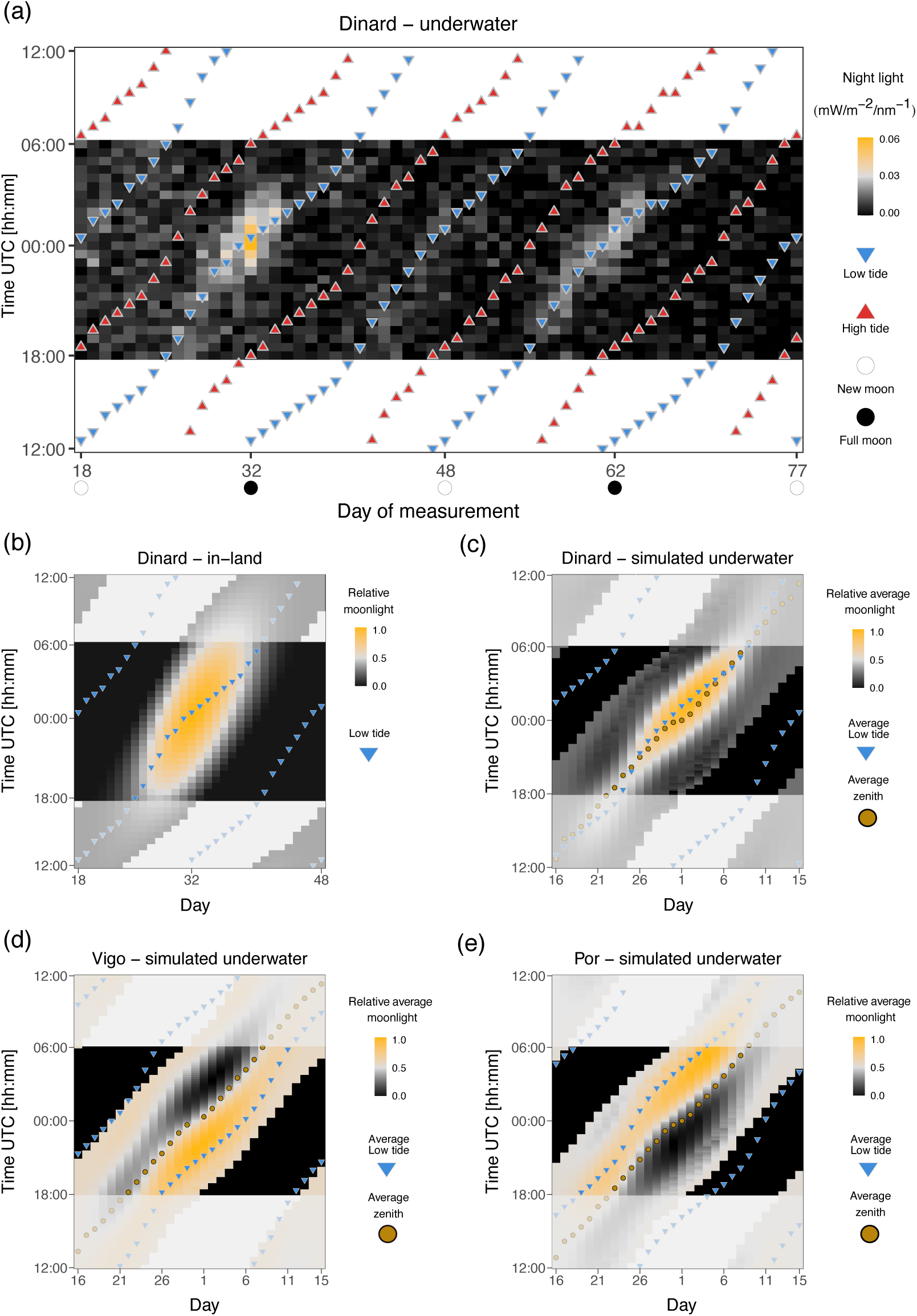
Moonlight intensity in the intertidal zone is modulated by the tides. (**a**) Heatmap of light intensity detected at night with a submerged radiometer in the intertidal region of Dinard. Days of two consecutive lunar months from new moon to new moon (2013-11-03 and 2014-01-02) are shown on the x-axis and time of day on the y axis. Timing of low and high tides is shown as blue and red triangles, respectively. The maximum daily moonlight duration detected was of 4h30 in days close to full moon. **(b)** Heatmap of relative moon illumination in-land for Dinard obtained from the R package ‘suncalc’. One month (from 2013-11-03 to 2013-12-04) is plotted and low tide is shown in blue inverted triangles. Night-time was considered to be the same as in the Dinard-underwater dataset, between 18h00 and 6h00. Moonlight duration in the absence of tides is over 14h on full moon days. **(c-e).** Heatmaps of relative average moonlight intensity for three geographic sites, simulated by multiplying normalized water levels and relative moonlight intensity (see Methods). Timing of low tides is shown in blue inverted triangles, moon zenith is shown in a yellow circle. **(c)** Dinard (France), **(d)** Vigo (Spain) and **(e)** Port-en-Bessin (France). Due to the effect of the water levels, the timing of moonlight visibility during the night varies greatly across geographical locations.

To assess the relative changes in light intensity over the day per wavelength, we used the raw 5-min light intensity dataset of 106 days (Figure S3a). Timing and water level of high and low tides were obtained from www.maree.info (Figure S3a) from the closest marine station available, located in St. Malo. The tidal timestamps were rounded to the nearest 5-minute interval to match the moonlight data.

### Tidal modulation of moonlight

In order to investigate how tides affect the visibility of moonlight in the intertidal region, we inferred the impact of the water column (Copernicus Program ERA5, see details below) on moonlight in-land (‘*suncalc*’ package, see below), and compared it with the quantified moonlight intensity underwater as measured by the radiometer (“Dinard-underwater” dataset).

### Comparing underwater and in-land moonlight patterns

We extracted water levels for Dinard (TableS1, FigureS1) from Copernicus Program ERA5 (Hersbach et al., 2020) for the same period of time as the “Dinard-underwater” dataset (Figure 1a). The ERA5 dataset comprises a reanalysis of water levels (a combination of model data with observations from across the world into a dataset using the laws of physics) with a temporal resolution of 10 min in UTC. Low tide and high tide were defined as the maximum and minimum water level in each tidal cycle, respectively (Figure 1a). In order to superimpose the 10-min water level dataset to the averaged 30-min light “Dinard-underwater” dataset, we assigned the tide timestamp to the closest 30-min interval (Figure 1a).

We also plotted moonlight intensity in-land, i.e. without tides, (R package *‘suncalc’,* Thieurmel and Elmarhraoui, 2022) at the same location over a single lunar cycle from new moon (03-11-2013) to new moon (04-12-2013) (Figure 1b). In further text we referred to this dataset as “Dinard in-land”. Relative moon illumination was calculated as the fraction of moon illumination in 10 min intervals multiplied by moon altitude. Low tides (blue triangles) were superimposed as above. Night-time was considered to be the same as in the “Dinard-underwater” dataset, i.e. between 18h00 and 6h00.

### Inferring light visibility in the intertidal of different geographical locations

The timing of the low tides varies across the Atlantic coast, and therefore tidal modulation of moonlight can be expected to differ in different geographic locations. As deploying the radiometer in different geographical locations was not feasible, we explored publicly available water and light datasets to simulate tidal modulation of moonlight. The comparison of field measurements from the intertidal zone (“Dinard-underwater”, Figure 1a) with general moonlight availability (“Dinard-in land”, Figure 1b) suggested that the tides considerably reduce the availability of moonlight. The simulations were set up in a way to reproduce this observed effect (compare Figure 1a to Figure 1c).

We extracted water levels (ERA5, Hersbach et al., 2020) and in-land moonlight intensities (R package *‘suncalc’,* Thieurmel and Elmarhraoui, 2022) in 10 min increments over eight lunar cycles from March to October of 2017, a time when *C. marinus* is not expected to be in diapause (Neumann, 1986).

In order to estimate how the water column affects moonlight intensity, we multiplied relative moonlight intensity with water levels scaled to a –1 to 1 range, referred as “simulated underwater” datasets. We then subsetted the data into eight cycles, where each cycle starts at midnight of a full moon day (full moon dates were taken from https://www.timeanddate.com). The eight cycles were averaged into one 30-day cycle (Figure 1c-e). Furthermore, we extracted the timing of the low tide as minimal water level value for each tidal cycle, subsetted the data into eight lunar cycles (day 1 = full moon), and averaged it (Figure 1c-e, blue triangles). Finally, we extracted the timing of the highest moonlight intensity (moon zenith) over the eight cycles, and averaged it accordingly (Figure 1c-e, golden circles). The analysis was performed for Dinard (“Dinard – simulated underwater”, Figure 1c), Vigo (“Vigo – simulated underwater”, Figure 1d) and Port-en-Bessin (“Por – simulated underwater”, Figure 1e). Night time was defined as 18h00 to 6h00, in order to allow the comparison between datasets.

### *Clunio marinus* laboratory culture

Two *C. marinus* laboratory strains were used in the moonlight entrainment experiments. Por-1SL strain (sampled in Port-en-Bessin, France, Figure S1) emerges during both full moon and new moon, therefore called semilunar (SL, see (Kaiser et al., 2021) for nomenclature of timing strains) and Vigo-2NM strain (sampled in Vigo, Spain, Figure S1) which emerges every new moon (NM). The strains were reared according to (Neumann, 1966), at 18°C (± 1.5°C) and under a light-dark (LD) cycle of 16:8 hours with a 4000 K neutral white LED light (Hera 61001491201) in all experiments. The moonlight provided varied depending on the experiment (see below).

For setting up experimental boxes, fertilized egg clutches collected over four days from several culture boxes were de-jellied with 5% bleach for a minute and rinsed five times with pasteurized 50:50 seawater and deionized water. Around 250 eggs were placed in transparent plastic boxes (8 x 8 x 8 cm) with 160 ml of pasteurized water (50:50 seawater and deionized water) and 2g of sand collected in different natural habitats of *C. marinus* (France and Portugal). Larvae were fed twice a week with 2ml of diatoms (*Phaeodactylum tricornutum*, strain UTEX 646). The water was exchanged every two weeks and after that the larvae were supplied with powdered nettles (*Urtica sp.*, Phoenix, Pharmahandel GmbH & Co KG). For one experiment, the simulated natural moon light with 6h high intensity, the boxes were set up differently: 20 fertilized egg clutches (around 1000 eggs) were washed in deionized water for one hour and placed in larger plastic boxes (20 x 20 x 5 cm). Larvae were fed twice a week with 5ml of diatoms and the water was exchanged as mentioned above.

Emergence phenotypes were obtained by counting the number of adults every day in the morning or early afternoon, over a minimum of two moonlight cycles and in three to six replicate boxes. Both strains emerge typically in late afternoon or night, so that the emergence day was recorded as the day before collection. The total number of individuals per cycle per treatment and replicate is given in TableS2. The fraction of emerged midges per experiment is given in Table S3.

Experimental boxes were placed in acclimatized rooms or in a I36-LL Percival incubator (Percival Scientific, USA). Light and temperature were measured over the course of the experiments with HOBO Pendant data loggers (UA-002-08 or UA-002-64, Onset Computer Corp, USA). A Quantum PAR Radiometer (Irradian Ltd., Scotland, UK) was used to characterize in detail irradiance (W/m2) and illuminance (lux) of the moonlights tested in this study (TableS4).

## Moonlight entrainment experiments

### Experiment 1: Moonlight intensity

In order to test if moonlight intensity affects the strength or phase of entrainment, we entrained the strains with different light intensities. Moonlight was given throughout the night for four consecutive nights every 30 days in a parallelized set up (Figure 2). Day 1 in the moonlight cycle was considered to be the first day the strains were subjected to light at night. Moonlight was simulated using white LED Array lights (LIUCWHA from Thorlabs, USA) dimmed with different neutral density filters (NDUV10B, NDUV20B and NDUV30B, Thorlabs, USA) to consistently decrease light intensity: 1) NDUV10B nominal optical density (OD) of 1, 2) NDUV30B with OD3 and 3) NDUV20B and NDUV30B in combination to obtain OD5. Irradiance and illuminance (TableS4) were measured using a ILT950 spectroradiometer (International Light Technologies, Peabody, MA, USA). Depending on the position in the acclimatized room, the light intensities were 40-200 lux for OD1, 0.4-2 lux for OD3 and 0.004-0.02 lux for OD5. In nature, moonlight intensity during full moon is expected to be between 0.05 and 0.2 lux, i.e. in between treatments OD3 and OD5. As these intensity values were not obtained from measurements underwater, it might be that light intensity reaching the developing larvae is lower than these ranges.

**Figure 2.**
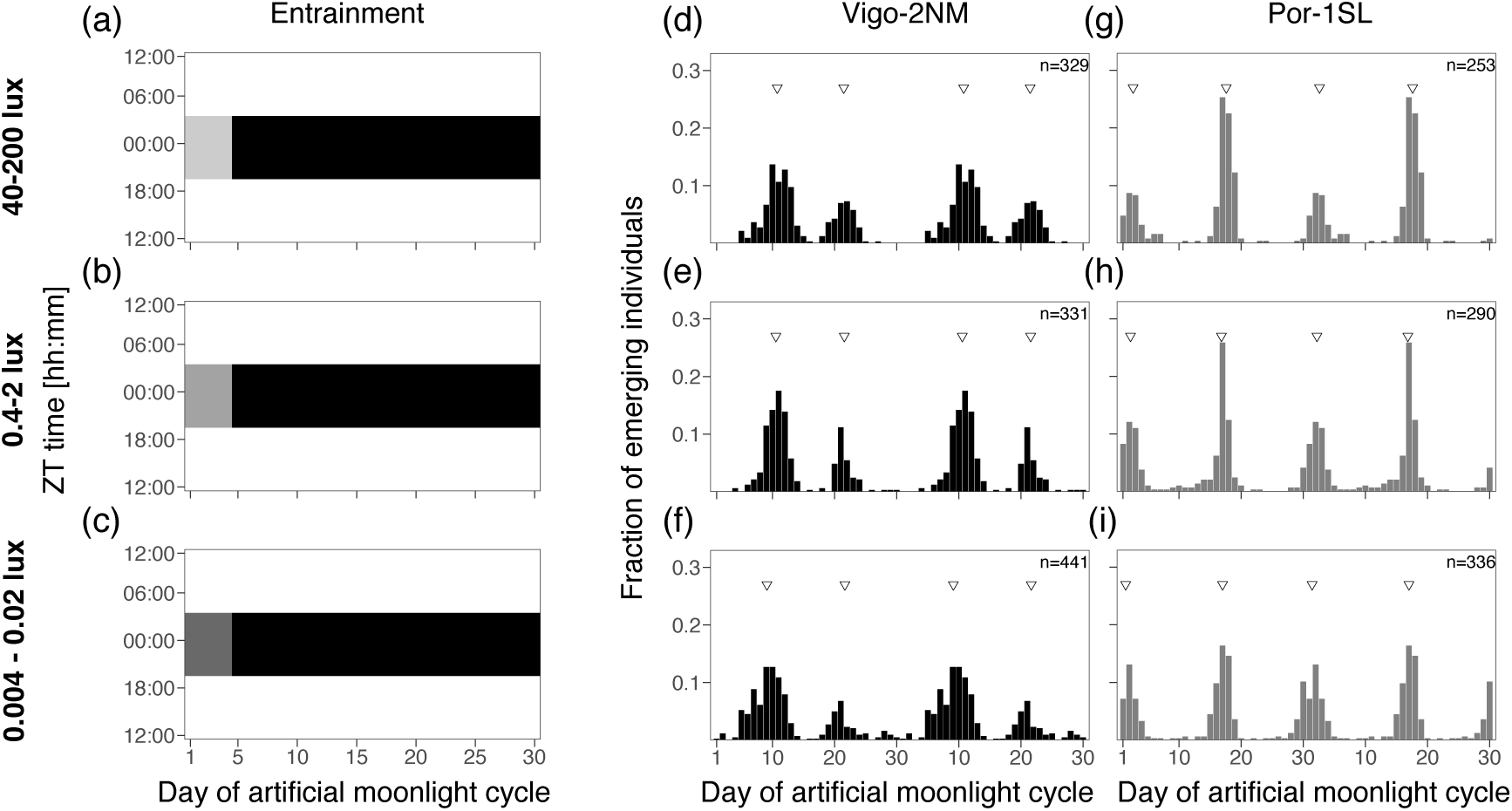
Moonlight intensity does not set the phase of the semilunar or lunar rhythm. (**a-c**) Artificial moonlight was provided throughout the night (8 hours) for 4 consecutive nights in a 30-day artificial moonlight cycle. Three light intensities were tested **(a)** 40-200 lux, **(b)** 0.4-2 lux and **(c)** 0.004-0.02 lux (TableS4). Heatmaps illustrate the moonlight treatment, where the x-axis represents days of moonlight cycle and y-axis shows zeitgeber time (ZT). Day 1 of the moonlight cycle is considered the first day moonlight is provided. Daylight is depicted in white, night in black and the artificial moonlight in different shades of grey. Emergence of the strains Vigo-2NM **(d-f)** and Por-1SL **(g-i)** are shown in barplots. Fraction of emerging individuals (y-axis) is shown over the moonlight cycle (x-axis), double-plotted for visualization. Total number of individuals per strain (values on top right) correspond to several replicate boxes over consecutive cycles summed up (TableS2). Inverted triangles show the phase of the rhythm detected with CircMLE (see Methods, TableS5). Differences in phase of entrainment are marginal across treatments for both strains (compare triangles).

### Experiment 2a and 2b: Two-hour and four-hour sensitivity windows

In order to test if there are specific windows of light at night for which the two *C. marinus* strains are exclusively sensitive to moonlight, windows of two or four hours over four consecutive nights were provided every 30 days. Day 1 of the moonlight cycle was considered as above. Zeitgeber Time (ZT) 0 is defined as the middle of their subjective night. For the two-hour window experiment (Figure 3), four discrete windows were tested over the eight hours of darkness: ZT 20-22, ZT 22-24, ZT 24-2 and ZT 2-4. Moonlight was mimicked with a 4000 K neutral white LED light (VT-2216, 7492, Pollin Electronic GmbH, Germany), with light intensity around 325 lux (TableS4). For the four-hour window experiment (Figure 4), three overlapping windows were tested: ZT 20-24, ZT 22-2 and ZT 0-4. Moonlight was provided with a White LED Array lights (LIUCWHA from Thorlabs, USA) with a NDUV10B filter and light intensity 40-200 lux (TableS4).

**Figure 3.**
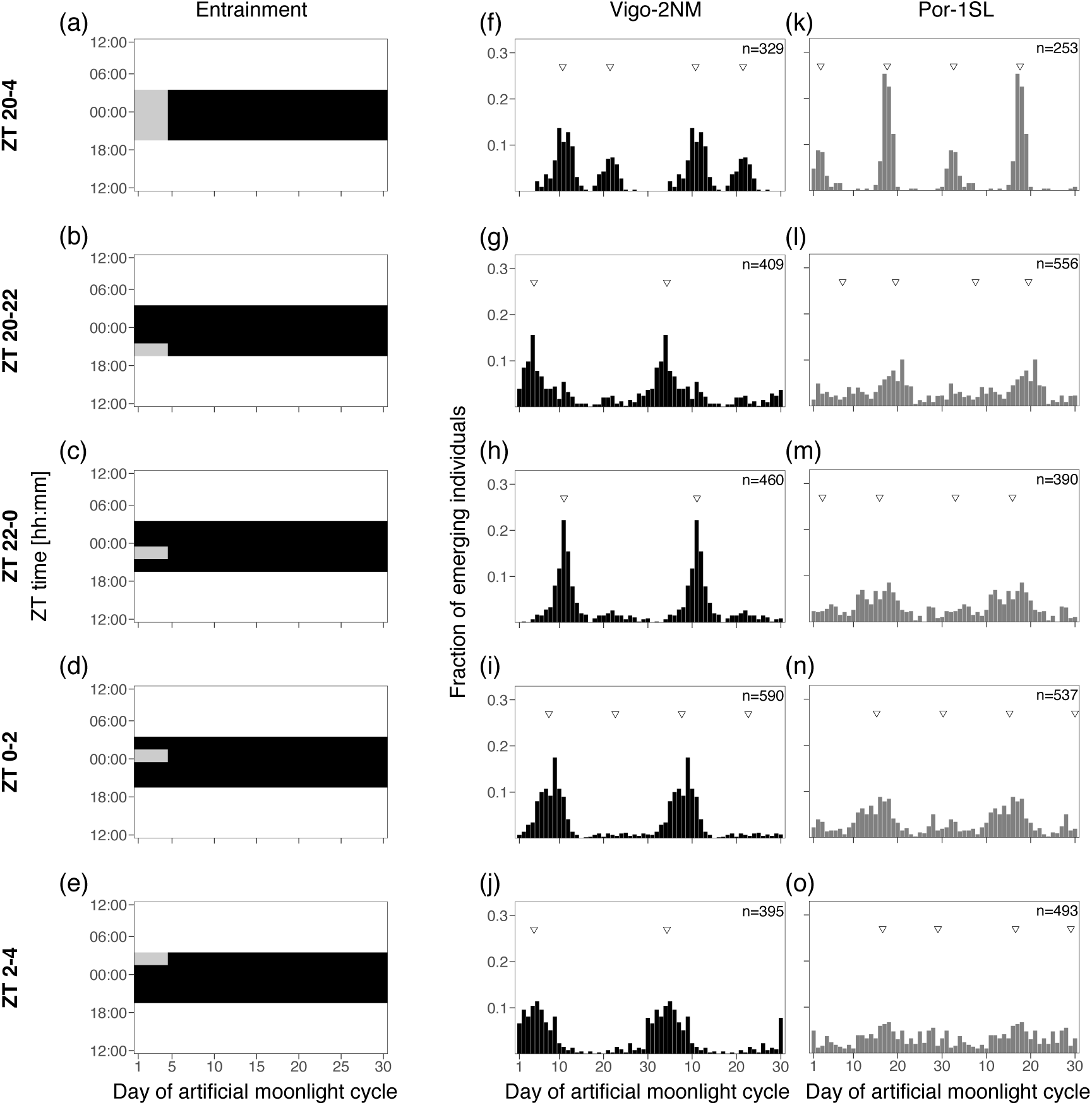
Light at night in two-hour windows entrains the rhythm in the Vigo-2NM strain but not the Por-1SL strain. (**a-e**) A block of 8h of artificial moonlight (standard treatment) was compared with 2h of artificial moonlight over four days at different times of the night (light grey rectangles). **(a)** Standard treatment of 8h of light at night (data from Figure 2), **(b)** moonlight from ZT 20-22, **(c)** moonlight from ZT 22-0, **(d)** moonlight from ZT 0-2 and **(e)** moonlight from ZT 2-4. Heatmaps illustrate the different entrainments tested, where the x-axis shows days of moonlight cycle and y-axis shows ZT time (ZT 0 is the middle of the night). Emergence of the strains **(f-j)** Vigo-2NM and (k-o) Por-1SL. Barplots show the fraction of emerging individuals (y-axis) over the moonlight cycle (x-axis), double-plotted for visualization. Total number of individuals per strain (values on top right) correspond to several replicate boxes over consecutive cycles summed up (TableS2). Inverted triangles show the detected phase of the rhythm with CircMLE (see Methods, TableS5). In Vigo-2NM strain, strength of entrainment varies depending on the timing of the moonlight at night and is highest in the window ZT 20-0 (note that the artificial peak is absent). In this window, the phase of the rhythm matches the standard treatment shown in (f, data from Figure 2d). The later the light is given at night, the earlier emergence occurs (TableS5). In contrast, none of the 2h-windows properly entrain the circasemilunar clock of Por-1SL strain (compare with k, data from Figure 2d, TableS5).

**Figure 4.**
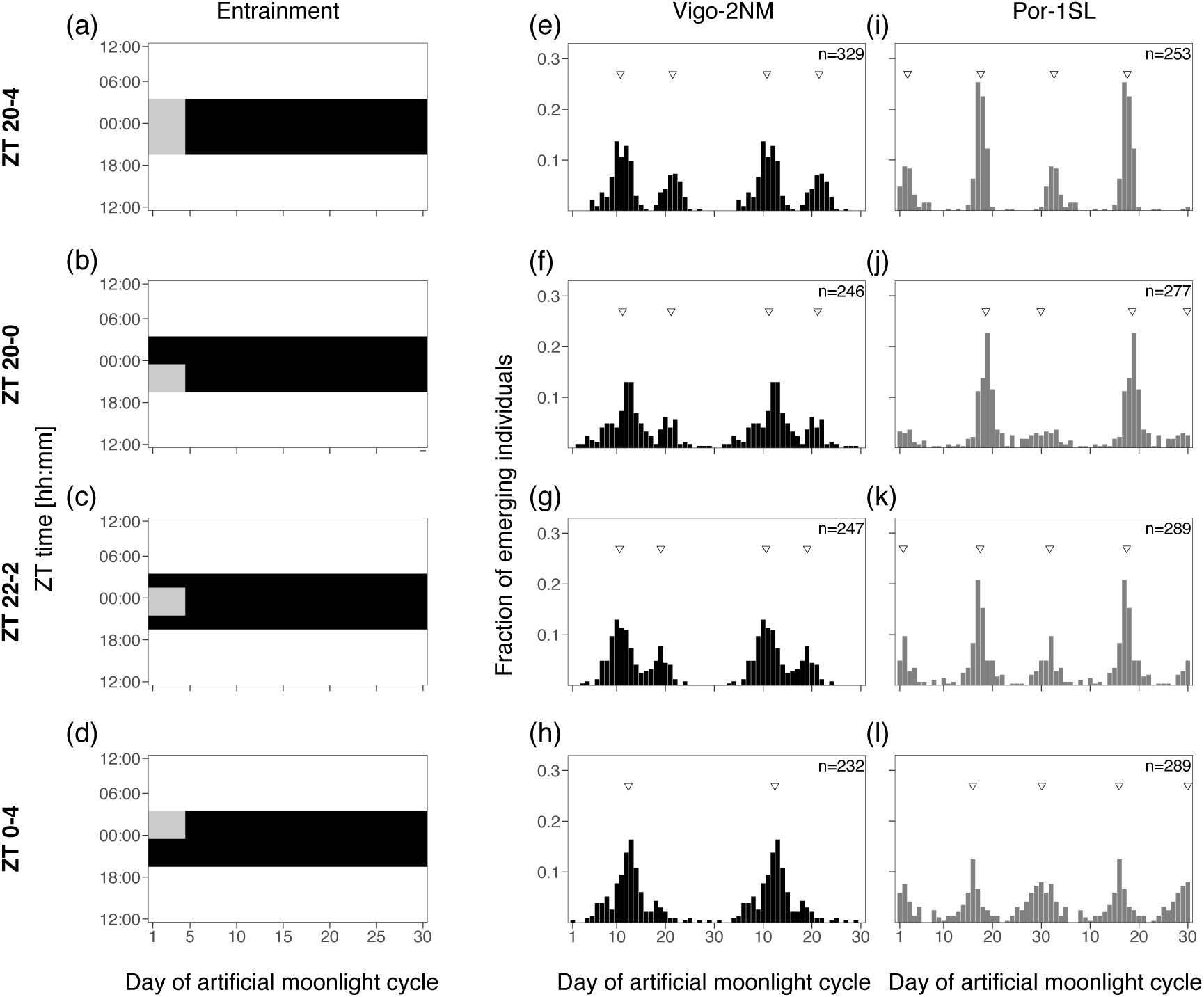
Light at night in four-hour windows entrains the rhythm in both strains. (**a-e**) A block of 8h of artificial moonlight (standard treatment, Figure 2a) was compared with 4h of artificial moonlight over four days at different times of the night (light grey rectangles). **(a)** Standard treatment of 8h of light at night (data from Figure 2a), **(b)** moonlight from ZT 20-0, **(c)** moonlight from ZT 22-2 and (d) moonlight from ZT 0-4. Heatmaps illustrate the different entrainments tested, where the x-axis shows days of moonlight cycle and y-axis shows ZT time (ZT 0 is the middle of the night). Emergence of the strains **(e-h)** Vigo-2NM and **(i-l)** Por-1SL is shown. Barplots show the fraction of emerging individuals (y-axis) over the moonlight cycle (x-axis), double-plotted for visualization. Total number of individuals per strain (values on top right) correspond to several replicate boxes over consecutive cycles summed up (TableS2). Inverted triangles show the detected phase of the rhythm with CircMLE (see Methods, TableS5). All 4h windows entrain Vigo-2NM and Por-1SL strains but strength of entrainment is lower in Vigo-2NM (compare with standard treatment and 2h-window ZT 22-0 in Figure 3h). In Por-1SL strain, strength of entrainment is high for all windows (TableS5). In Vigo-2NM strain, the windows ZT 20-0 and ZT 22-2 match the phase of the standard treatment whereas in Por-1SL strain the match is only observed for window ZT 22-2.

### Experiment 3: Shifting block of moonlight

In *C. marinus* natural habitat, moonlight intensity is a complex environmental cue that depends on the lunar phase and on the tidal cycle. Every day, moonrise and moonset shift by about 48 min. Likewise, tidal cycles have a period of 12.4h and high tide and low tide shift every day by about 48 min and can hinder moonlight visibility (see above and discussion). Here, we tested if the natural timing of moonlight alone (without intensity changes) affects the entrainment of the circalunar clock. To this end, we simulated moonlight as a block of light (two, four or six hours duration) shifting every day by 48 min. This resulted in a cycle of 31 days with 30 moonlights (Figure 5). Moonlight was simulated using white LED Array lights (LIUCWHA from Thorlabs, USA) together with a neutral density filter NDUV10B (intensity 40-200 lux, TableS4) connected to an Arduino Mega 2560 microcontroller (Arduino, Italy, http://www.arduino.cc/). Custom-made scripts were written to turn the lights on and off with the duration of a lunar-day, i.e. with a daily shift of 48 min. The intrinsic error rate of each Arduino was assessed by measuring the duration of light transitions over consecutive days with HOBO Pendant (measurements each 10s, data not shown). This error was then taken into consideration in each loop run by the Arduino in order to obtain the expected timings. To match the emergence patterns across treatments, we considered day 1 as the day when the middle of the moonlight treatments occurs at ZT0 (asterisks in Figure 5).

**Figure 5.**
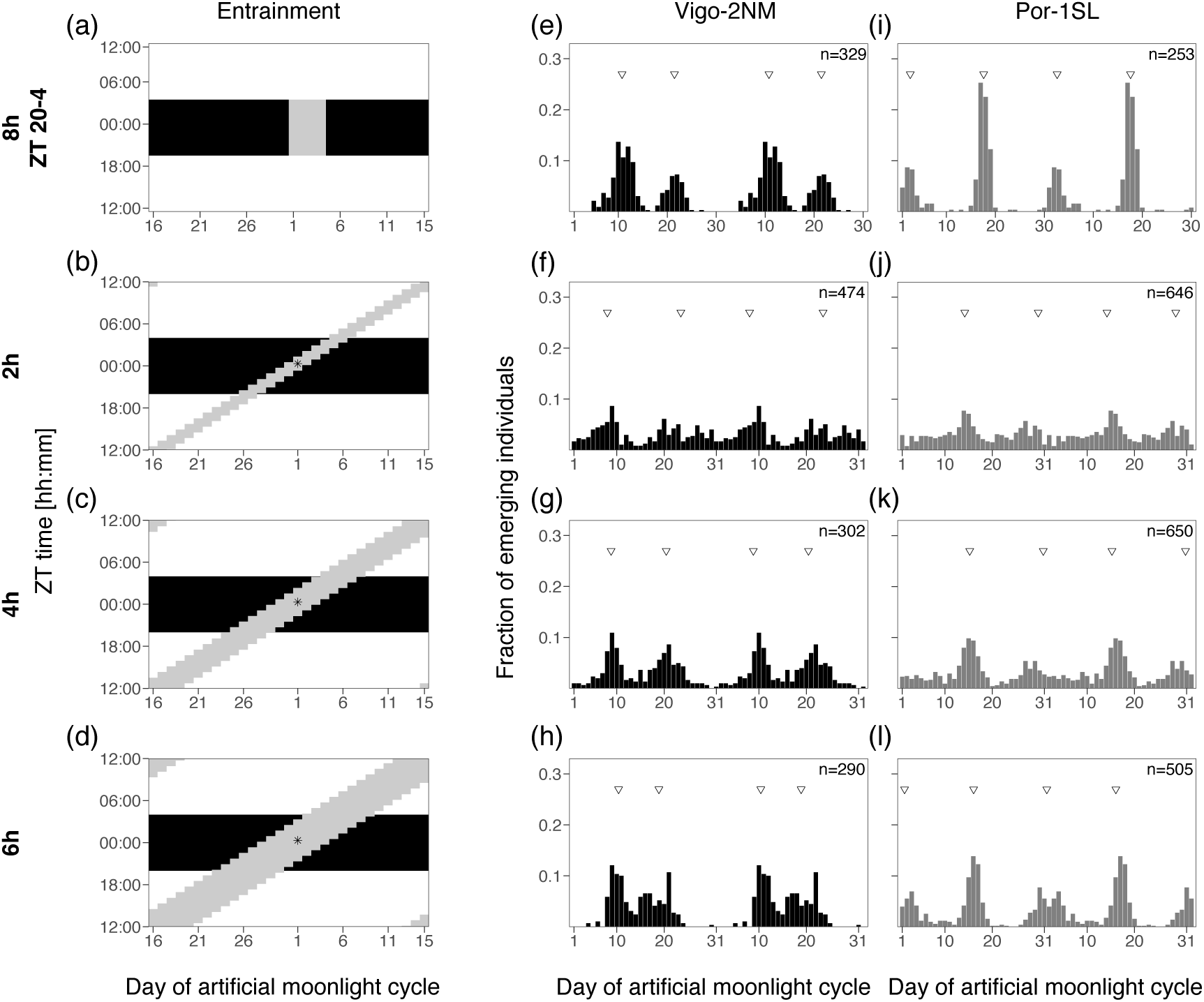
Duration of a shifting-block of moonlight impacts the strength of entrainment the circa(semi)lunar clock. (**a-d**) A block of 8h of artificial moonlight (standard treatment, Figure 2a) was compared with moonlight presented in a lunar day cycle (24.8 h), i.e. with a daily shift of 48 min of a high-intensity moonlight (40-200 lux, light grey shading). The daily shift of 48 min results in a moonlight cycle of 31 days (x-axis). Y-axis shows ZT time. Day 1 in all entrainments was considered the day where the middle of the moonlight occurs at ZT 0, the middle of the night (asterisk). Moonlight duration was **(b)** 2 hours, **(c)** 4 hours or **(d)** 6 hours. Emergence of the strains **(e-h)** Vigo-2NM and **(i-l)** Por-1SL. Barplots show the fraction of emerging individuals (y-axis) over the moonlight cycle (x-axis), double-plotted for visualization. Total number of individuals per strain (values on top right) correspond to several replicate boxes over consecutive cycles summed up (TableS2). Inverted triangles show the detected phase of the rhythm with CircMLE (see Methods, TableS5). The 2h shifting-block is insufficient to properly entrain the circa(semi)lunar clock of Por-1SL or Vigo-2NM strains. The phase of the rhythm under 6h and 4h shifting-blocks matches the standard treatment in both strains. This suggests that the strains are most sensitive to moonlight in the middle of the night.

### Experiment 4: Simulated natural moonlight

In this experiment, we simulated both the natural timing of moonlight (i.e. the daily shift), as well as changing moonlight intensity over the night and over the lunar cycle. This was achieved with a Profilux Light computer (GHL, Germany) with a customized firmware. Briefly, the light program consists of a sine wave of light intensity applied to a 30-day cycle to simulate lunar phases. Then, a second sine wave of light intensity is applied to the 24 hours light cycle, simulating the rise and fall of the moon or the effect of water levels respectively. The day with maximum moonlight intensity across the lunar cycle is defined as full moon and on that day, the daily maximum moonlight intensity is set to the middle of the night (asterisks in Figure 6). This was considered as day 1 of the moonlight cycle. Maximum light intensity shifts every day for 48 min simulating the 24.8h lunar day. The width of the daily sine-wave (four hours or six hours) corresponds to time length of light provided to further simulate the effect of the water levels on light visibility. We applied three different treatments: 1) four-hour moonlight of low intensity (average maximum intensity around 0.26 lux, Table S4), 2) six-hour moonlight of low intensity (average maximum intensity around 0.24 lux, Table S4) and 3) six-hour moonlight of high intensity (average maximum intensity around 900 lux, Table S4). Because the program calculates light intensities per day and fits the 48-minute shift to a 30-day cycle, there is a jump in light intensity at midnight for the 6-hour high intensity moonlight treatment (see Figure 6j and Figure S4a). For the other treatments, the recalculation was set to occur during the day and thus moonlight during the night was not affected (Figure S4b). The computer was connected to a dimmable (by pulse width modulation) LED Mitras Lightbar 2 Actinic 120 (PL-1294, GHL, Germany), using cold white (8000 K) color channel and a maximum light intensity set to 100%. For treatments 1) and 2), light intensity was reduced by covering the lightbar with aluminium foil, punctured evenly over its length. The lightbars were placed at the back of I36-LL Percival incubators (Percival Scientific, USA) and irradiance and illuminance (TableS4) were measured using a ILT950 spectroradiometer (International Light Technologies, Peabody, MA, USA) to ensure all replicate boxes were exposed to the same light intensity.

**Figure 6.**
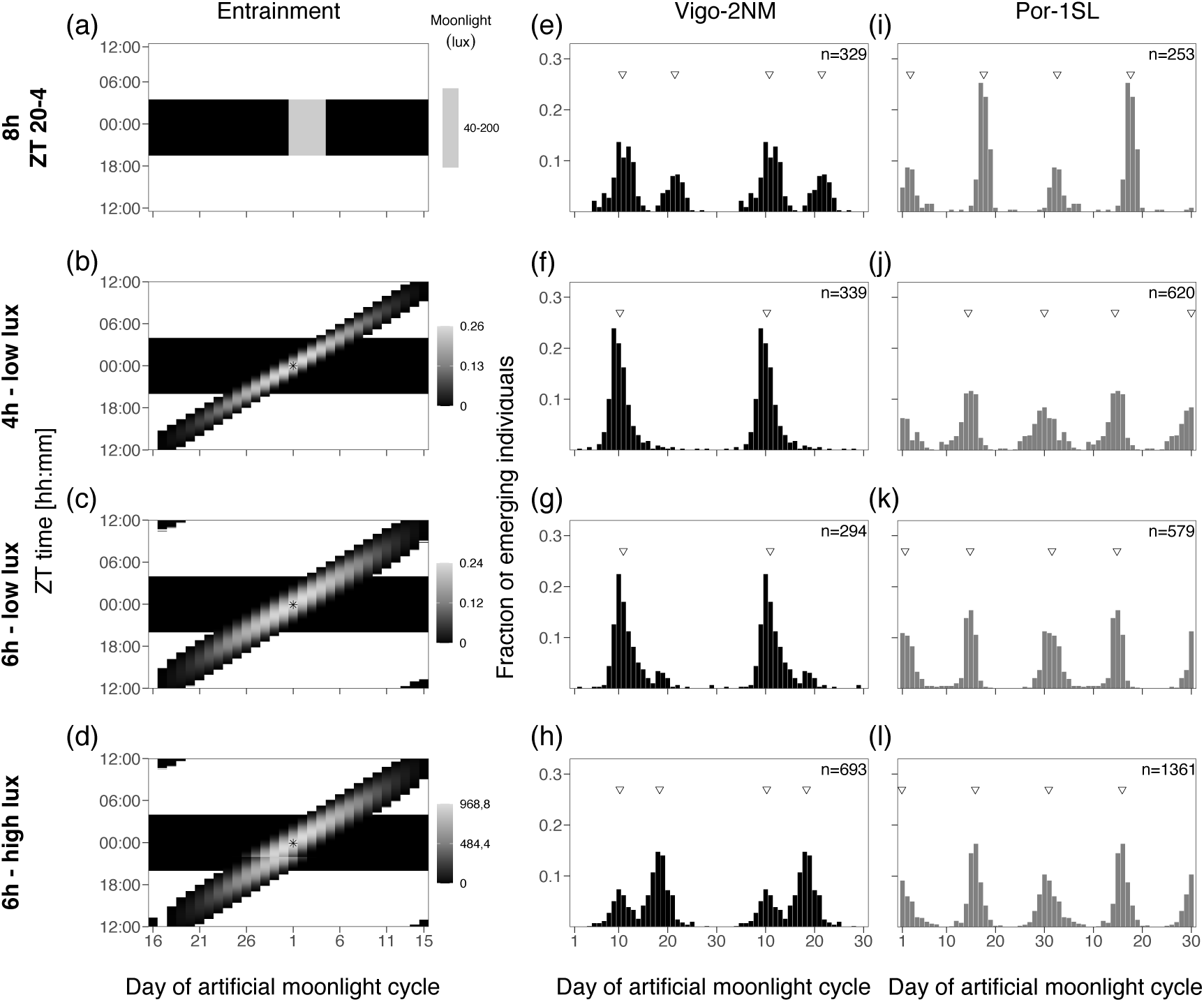
Light intensity in simulated natural moonlight impacts strength but not phase of entrainment. (**a-d**) A block of 8h of artificial moonlight (standard treatment, Figure 2a) was compared with moonlight presented in a lunar day cycle (24.8 h), i.e. with a daily shift of 48 min together with light intensity modulation. Moonlight intensity was changing gradually over the night and over the lunar cycle. Heatmaps are coloured-scaled for light intensity throughout the 30-day moonlight cycle (x-axis) and ZT time (y-axis). Day 1 in all entrainments was considered the day where the middle of the moonlight occurs at ZT 0, the middle of the night (asterisk). Natural levels of moonlight during full moon (<0.3 lux) were tested with a duration of **(b)** 4 hours and **(c)** 6 hours. **(d)** High intensity moonlight (similar to the standard treatment, 600 lux) was tested with a duration of 6 hours. Emergence of the strains **(e-h)** Vigo-2NM and **(i-l)** Por-1SL. Barplots show the fraction of emerging individuals (y-axis) over the moonlight cycle (x-axis), double-plotted for visualization. Total number of individuals per strain (values on top right) correspond to several replicate boxes over consecutive cycles summed up (TableS5). Inverted triangles show the detected phase of the rhythm with CircMLE (see Methods, TableS2). In Vigo-2NM strain, both 4h and 6h low intensity light treatments entrain the circalunar clock without artificial peak. The 6h treatment of high intensity leads to a strong artificial peak that mirrors the observed emergence pattern in the 6h-shifting block. In Por-1SL strain all simulated moonlight treatments lead to a strong entrainment of the circasemilunar clock.

### Matching phase-setting moonlight windows in the laboratory and the field

We further explored how the phase-shifts detected in experiment 2a and 2b relate to the optimal adult emergence timings in the field. First, a “Vigo moon*tides 4h dataset” and a “Por moon*tides 4h dataset” were generated by filtering the respective “simulated underwater” dataset (see above; before averaging months) to allow the light to be visible for only 2h before and after low tide. The dataset was then subsetted into lunar cycles (day 1 = full moon) and averaged into one representative 30-day cycle (Figure 7a-b). The experimentally tested entrainment windows in experiments 2a and 2b were fitted into the moonlight windows of the “Vigo moon*tides 4h dataset” and “Por moon*tides 4h dataset” (coloured boxes in Figure 7a-b). For this fitting, the daylight timing in the field (Figure 7a-b, left y-axis) and the LD cycle from the experiment (Figure 7a-b, right y-axis) needed to be matched. To this end, we averaged sunrise and sunset intensity values (obtained from the R package ‘suncalc’) across the eight consecutive months in the two locations, leading to sunrise and sunset values to be set to the longest day. In Vigo, this resulted in a LD of 15:9 which was fitted in a LD 16:8, with the light phase between 4h30-20h30. In Port-en-Bessin the LD was 16:8 with the light phase being between 4h00-20h00. Based on this, the middle of the dark-phase was matched at ZT0 (Vigo at 00h30 UTC and Port-en-Bessin 00h00 UTC). Then each experimental 2h or 4h light window was placed on the field days for which there were at least two hours of moon visible in four consecutive days. This matching translates into a prediction of which days the observed emergence times in the laboratory would fall to in the field (Figure 7e-f). Finally, we compared these predictions to the optimal timing, i.e. the spring tide days. To this end, water levels during low tides (see section “Inferring light visibility in the intertidal of different geographical locations”) were averaged to indicate low tide levels (dots in Fig. 7c-d) and the corresponding spring tide days (the lowest low tide of the month +/− 2 days, grey shading in Figure 7 c-d and e-f).

**Figure 7.**
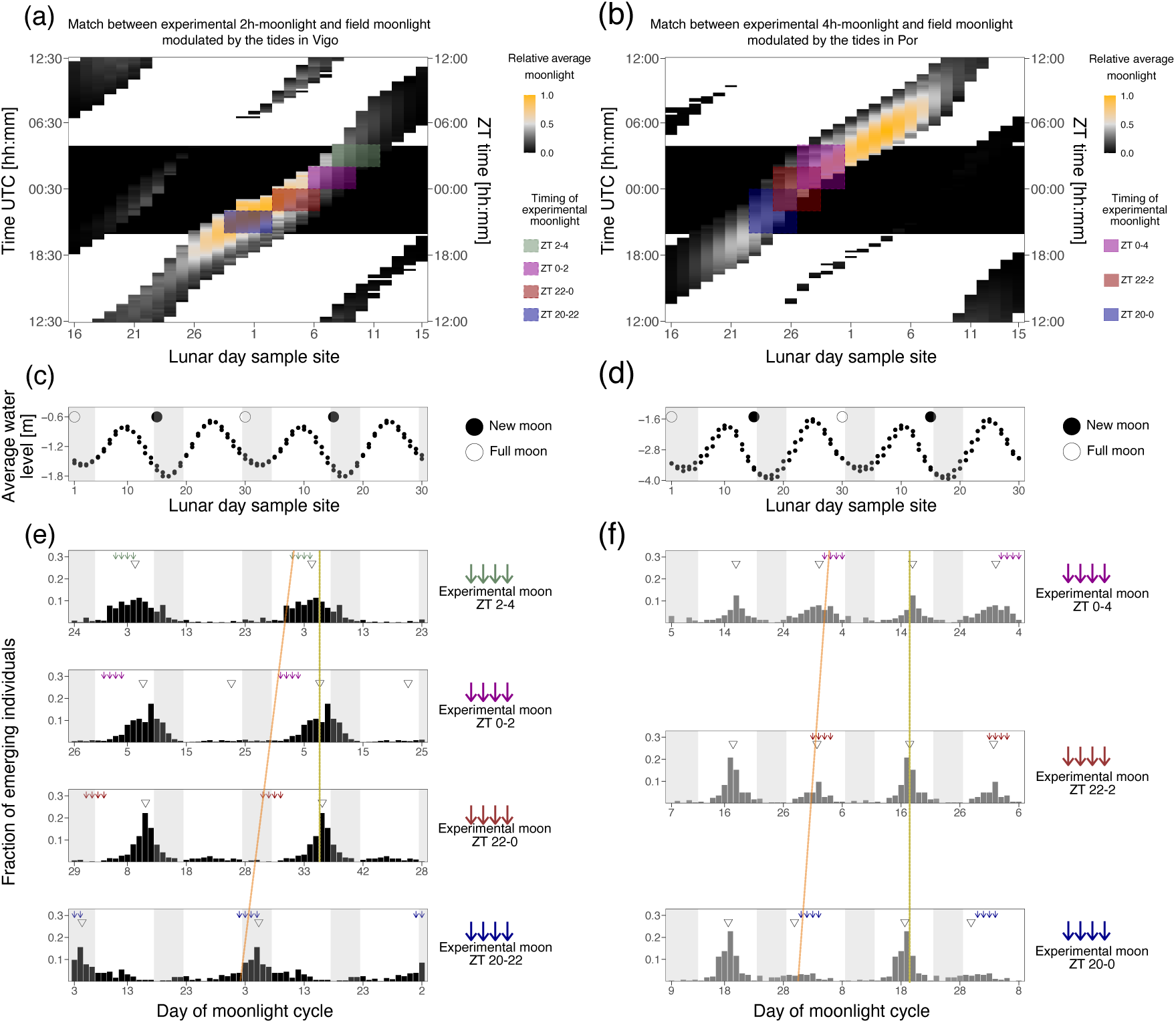
Phase-shifts due to timing of light at night might be adaptive and match the natural timing of emergence in Vigo-2NM strain, but not Por-1SL strain. (**a-b**) Heatmaps of simulated moonlight (water levels multiplied with relative moonlight intensity). Only four hours of light around low tides are shown. Daylight is shown in white and night in black. Days in the moonlight cycle are shown on the x-axis, the time in UTC is shown on the left y-axis and ZT time is shown on the right y-axis. The field light-dark cycle was fitted to a 16:8 LD cycle and matched with the 16:8 LD cycle from the experiment based on the middle of the night. Coloured boxes represent the tested sensitivity windows for **(a)** the Vigo-2NM strain under 2h moonlight windows and **(b)** the Por-1SL strain under 4h moonlight windows. The windows were matched to the field data so they had maximal overlap with the simulated moonlight for four consecutive nights. For the Vigo-2NM strain, this results in ZT 20-22 matching day 29-2 in field (blue rectangle), ZT 22-0 match day 3-6 (red rectangle), ZT 0-2 match day 6-9 (pink rectangle) and ZT 2-4 match day 8-11 (green rectangle). For the Por-1SL strain, ZT 20-0 matches day 23-26 in field (blue rectangle), ZT 22-2 matches day 25-28 (red rectangle) and ZT 0-4 matches day 27-30 (pink rectangle). **(c-d)** Averaged low tide water levels (y-axis) and respective times of new moon (black circle) and full moon (white circle) are shown in a 30-day cycle (x-axis; double-plotted for visualization). Grey shadings represent the four days with the lowest water levels, i.e. the spring tides. **(e)** Emergence of Vigo-2NM strain under 2h-windows at different times of the night. Emergence data from each tested window in Figure 3 was shifted according to when this window is expected to occur in the field (see panel a). Arrows represent days in which the moon was provided and are coloured by the 2h-window tested. Inverted triangles show the detected phase of the rhythm with CircMLE (see Methods, TableS5). Grey shadings represent the spring tides predicted in (c). Under most windows, emergence occurs just before the new moon spring tides, as is generally observed for new moon populations. Thus, the observed shifts in phase depending on the moonlight windows allow emergence to happen at the same time of the lunar cycle. Surprisingly, light at dusk leads to emergence in the spring tides of full moon. **(f)** Emergence of Por-1SL strain under 4h-windows at different times of the night. Again, the observed shift in phase of emergence ensures that emergence happens at the same time of the lunar cycle. However, emergence does not match the spring tide days, in contrast to what is observed in the field.

## Statistical analysis

### Phase and peak concentration from CircMLE

Phase and peak concentration were determined using the statistics software R (version 4.2.3) and the package CircMLE (Fitak and Johnsen, 2017). CircMLE uses a likelihood-based approach to analyse unimodal or bimodal circular data according to 10 defined models. Briefly, the uniform model (M1) has no significant directionality (arrhythmicity), unimodal models (M2A, M2B, M2C) have a single direction (single emergence peak) and bimodal models (M3A, M3B, M4A, M4B, M5A, M5B) have two significant directions on the circle (two emergence peaks). Each model is described by parameters such as mean direction (φ1 and φ2) and concentration (k1 and k2), which in chronobiology approximates the phase and peak concentration, respectively. We considered φ1 as the phase of the peak detected closer to day 1 (peak 1, TableS5) and φ2 as the phase of the second detected peak (peak2, TableS5). The likelihood that each dataset fits into each of the 10 models was evaluated, and the most likely model was selected according to the Akaike Information Criterion (AIC). AIC is the sum of the model fit (deviance) and twice the number of parameters. It penalizes models with more parameters, thus the model with the lowest AIC was considered the most suitable (Table S5). We run CircMLE with no a priori assumption on the expected emergence distributions and considered all provided models to fully examine the effects of the light treatments.

### Rhythmicity index (RI)

In order to assess the strength of entrainment, we calculated the rhythmicity index (RI) for each experiment separately. RI was calculated according to (Levine et al., 2002) with a few modifications. In short, number of emerged individuals per day was first normalized per cycle (30 days) and autocorrelation was calculated using *acf* function of *tseries* R package v0.10-55 (Trapletti and Horni, 2023). RI represents the autocorrelation function of a lag of 16 for semi-lunar populations (period of 15 days) and of a lag of 31 for lunar populations (period of 30 days). Significance of the rhythm is defined as RI > 0.3 = highly significant, 0.1 < RI < 0.3 = significant, RI < 0.1 = not significant.

## Results

### Moonlight intensity is modulated by the tides

We measured light intensities in the intertidal zone of Dinard with a hyperspectral radiometer placed just below the mean low water level of spring tides. We found that the intensity of both moonlight and daylight is modulated by the tides (Figure 1a, Figure S2, Figure S3). Light intensities were the highest during low tide for both moonlight (Figure 1a) and daylight (Figure S3). The 50-min shift in the highest daily moonlight intensity following the 24.8h lunar-day can be observed, as well as the increasing and decreasing light intensity towards and after full moon days (Figure 1a). The highest intensities of both moonlight and daylight are found at wavelengths between 400 and 700 nm (Figure S2a and c). Normalizing daylight intensity for each wavelength category independently (Figure S2d) reveals that daylight is also detectable at wavelengths below 400 nm and above 700 nm and that tidal modulation of light is relatively uniform over the measured spectrum (Figure S3). However, it is not possible to distinguish the signal of moonlight from background noise at these wavelengths (Figure S2a-b). This shows that moonlight can only be effectively detected by the radiometer in the range of 400-700 nm.

Daylight was detected during every low tide with similar intensities (Figure S2c). In contrast, moonlight is primarily detected during the night-time low tides around full moon (Figure S2a). During new moon there are also traces of light at night, but much less intense than during full moon (Figure S2a), suggesting that the low ambient light during new moon is still modulated by the tides.

In order to explore by how much the water levels restrict the availability of moonlight, we compared the light measurements we obtained from the intertidal zone (Figure 1a), with calculated moonlight intensities in-land (Figure 1b). While relative moonlight illumination over Dinard during full moon can last up to 14 hours (Figure 1b), in the measurements from the intertidal zone it is only detectable above background noise for approximately 4-5 hours (Figure 1a).

In each geographical location, the timing of the tides varies. Consequently, the timing of light at night and the lunar phase during which highest light intensities reaching *C. marinus* larvae may differ. We tested if combining the publicly available datasets of in-land moonlight and water levels in Dinard would recapitulate the moonlight patterns observed in underwater measurements. There is a good correspondence between the calculated patterns (Figure 1c) and the field measurements (Figure 1a). Therefore, we used this simulation to predict the patterns of moonlight visibility at the sites where *C. marinus* strains used in entrainment experiments described below were sampled (Figure 1d-e). In Dinard, the days with low tides in the middle of the night correspond to full moon days (Figure 1c). However, this is not the case for other locations along the coast. Therefore, the patterns differ from those found in Dinard in three aspects (compare Figure1c-e). First, moonlight available in the middle of the night occurs several days before full moon (Por; Figure 1e) or after full moon (Vigo; Figure 1d). Second, the highest moonlight intensity around full moon is late in the night (Por; Figure 1e) or early in the night (Vigo; Figure 1d). Third, some low-intensity moonlight may also be visible in a second window just before new moon (Por; Figure 1e) or just after new moon (Vigo; Figure 1d).

In entrainment experiments, which are described below, we tested how intensity, duration and time of light at night affect the strength and phase of entrainment and how adult emergence matches the timing of lowest tides based on the prediction when *C. marinus* strains would perceive light at night in their respective geographical location.

### Experiment 1: Moonlight intensity does not set the phase of the semilunar or lunar rhythm

Four consecutive nights with dim illumination were previously established as the standard artificial moonlight treatment for entrainment of *C. marinus’* circalunar clock (Neumann, 1966).

Here, we first asked if exposing the strains Vigo-2NM and Por-1SL to different moonlight intensities under the standard treatment would affect the strength or phase of entrainment. A light set up with optical density filters was used to provide three independent treatments, namely 40-200 lux (Figure 2a), 0.4-2 lux (Figure 2b) and 0.004-0.02 lux (Figure 2c). We found that phase of entrainment is unaffected by light intensity in both Vigo-2NM (Figure 2d-f) and Por-1SL (Figure 2g-i) strains. Across treatments, both peak 1 and peak 2 differ in phase by less than a day (TableS5). Notably, lower light intensities do not reduce the strength of entrainment in Vigo-2NM (Figure 2f) and Por-1SL (Figure 2i) strains, as observed in the Rhythmicity Index RI (Table S5). This confirms that *C. marinus* is exquisitely sensitive to moonlight down to at least 0.004 lux.

In all treatments, the Vigo-2NM strain shows a smaller second emergence peak, about 10 days after the first peak and not fitting a semi-lunar pattern. The second peak is stronger at higher light intensities (Figure 2d vs Figure 2f), which is reflected in an increasing concentration of the second peak with increasing light intensities (see k2, Table S5) and in the absence of a significant rhythmicity index at the highest light intensity (RI = 0.02). This peak is considered an artifact of the highly artificial moonlight regime, as it is absent in the field and also under more realistic moonlight regimes in the laboratory (Kaiser et al., 2021). The experiments below also explore which moonlight regimes lead to an emergence pattern without the artificial peak.

### Experiment 2a: Two hours of light in the middle of the night set the phase of the rhythm in the Vigo-2NM strain but not in the Por-1SL strain

To explore (1) if a moonlight sensitivity window as described for the San strain (Neumann 1969) also exists in Por-1SL and Vigo-2NM strains, if (2) its duration and/or timing in relation to the light-dark cycle is strain-specific, and if (3) it sets the phase of the circalunar clock, we exposed these strains to a bright artificial moonlight in two-hour windows. We applied four independent treatments consisting of windows corresponding to “after dusk” (ZT 20-22, Figure 3b), “middle of the night” (ZT 22-0, Figure 3c and ZT 0-2, Figure 3d) and “before dawn” (ZT 2-4, Figure 3e). We compared emergence patterns under these entrainments with the emergence under the standard treatment with eight hours of light at night (Figure 3a, f, k), replotted from Figure 2a, d, g). We found that two hours of light at night entrain the circalunar clock of the circalunar strain Vigo-2NM. Strength, rhythmicity and phase of entrainment differ depending on the timing of light at night (Figure 3g-j). Shorter moonlight durations clearly reduce the strength of the artificial peak. Except for the window ZT 0-2, where the model that best describes the emergence distribution is bimodal (M4B, TableS5), unimodal distributions were detected (as expected for a circalunar strain).

Depending on the timing of moonlight, the strength of entrainment varies greatly (see RI, TableS5). The window ZT 22-0 (Figure 3h) not only leads to the strongest peak concentration and highest RI (TableS5), but also the phase of the emergence patterns matches the standard treatment (compare Figure 3h with Figure 3f). This suggests that Vigo-2NM strain is most sensitive to moonlight during the two hours prior to the middle of the night. Providing light at night two hours after the middle of the night (ZT 0-2) leads to an advance in the phase of emergence of around three days compared to the standard treatment. Interestingly, moonlight windows after dusk (ZT 20-22; Figure 3g) and before dawn (ZT 2-4; Figure 3j) windows lead to a phase advance of around six days, but strength of entrainment is low (see RI, TableS5).

In contrast, for Por-1SL strain we found that two hours of light at night do not properly entrain the circasemilunar clock (Figure 3l-o). In all treatments, there are individuals emerging throughout the lunar cycle, which is not observed in the standard treatment (compare Figure 3k with Figure 3l-o). CircMLE detects bimodal distributions for all treatments but with low concentration peaks (on average less than 2, TableS5) and only the window from ZT 0-2 had a significant RI (TableS5).

### Experiment 2b: Four hours of light in the middle of the night entrain the rhythm in both strains

As Por-1SL strain was not entrained by two hours of light at night, we tested if increasing the duration of light to four hours would entrain its circasemilunar clock. Alongside, we also tested this treatment in Vigo-2NM strain. We used three different windows of light at night, ZT 20-0 (“after dusk”, Figure 4b), ZT 22-2 (“middle of the night”, Figure 4c) and ZT 0-4 (“before dawn”, Figure 4d), and compared to the standard treatment (Figure 4a, e, i, replotted from from Figure 2a, d, g). We found that in Por-1SL strain, four hours of light at night are sufficient to entrain the circasemilunar clock (Figure 4 j-l). In all treatments, emergence displays a bimodal distribution (M5B, TableS5). There are some differences in the concentration of peaks across treatments, with the moonlight window in the middle of the night (ZT 22-2, Figure 4k) leading to the strongest rhythmicity (RI=0.35, Table S5). The emergence distribution under this treatment also most closely matches the standard treatment (compare Figure 4k with Figure 4i). The phase of emergence changes slightly with the timing of the light window (Figure 4j-l; Table S5). Taken together, these results suggest that the Por-1SL strain requires longer moonlight durations than the Vigo-2NM strain and is most sensitive to moonlight in the middle of the night.

In Vigo-2NM strain, four hours of light at night lead to less concentrated peaks in most treatments compared to two hours of light at night (Figure 4f-h compared with Figure 3h). For the 4-hour moonlight windows most models suggest a bimodal distribution, showing that this entrainment is less optimal for Vigo-2NM strain as it increases the artificial peak around day 20 (see k2, Table S5).

The different treatments entrain the rhythm in slightly differing phases (–1 to –2 days, see peak 1 in Table S5). The window in the middle of the night (ZT 22-2, Figure 4g) sets the phase similarly to the standard treatment. The window after dusk (ZT 20-0, Figure 4f) induces a delay of less than a day while the before dawn window leads to a phase delay of almost two days (ZT 0-4, Figure h) with a strong strength of entrainment (RI = 0.38, Table S5).

### Experiment 3: Duration of a shifting-block of moonlight impacts the strength of entrainment of the circa(semi)lunar clock

With the simplified artificial moonlight regimes tested up to here, we were able to show that 1) moonlight intensity does not affect the phase or strength of entrainment, but induces an artificial peak in Vigo-2NM strain, 2) duration of light at night required to entrain the rhythm is strain-specific, 3) light presented in the middle of the night sets the phase similar to the standard moonlight treatment, and 4) the timing of light at night usually shifts the phase of the lunar rhythm. However, moonlight is a complex environmental cue, with intensities changing across the lunar cycle and due to the effect of the tides (Figure1). Therefore, we next tested a shifting-block moonlight treatment which simulates the daily shift of the moonlight onset and offset of 48 min under a 31-day cycle with a high intensity moonlight (40-200 lux, as the standard treatment). The effect of the tides hindering moonlight visibility was simulated by changing the duration of the of block of light: two hours (Figure 5b), four hours (Figure 5c) or six hours (Figure 5d).

The two-hour shifting block of moonlight neither entrained the circalunar clock in Vigo-2NM strain (Figure 5f) nor the circasemilunar clock in Por-1SL strain (Figure 5j), even though in Vigo-2NM a non-shifting two-hour window of moonlight entrained the clock effectively (Figure 3g-j). The four– and six-hours shifting-block treatments lead to reasonable entrainment in Vigo-2NM strain (Figure 5g and h, respectively) and Por-1SL strain (Figure 5k and l, respectively). However, in Vigo-2NM strain the artificial peak is particularly pronounced and bimodal distributions were detected (Table S5), suggesting that as in experiment 2b the longer moonlight (4h and 6h) increases the artificial peak.

In the shifting-block treatments, we defined day 1 as the day that the middle of the moon hits the middle of the night (Figure 5b-d, asterisk). Under this definition of day 1, the phase of emergence in the shifting-block moonlight and in the standard treatment (Figure 5a, e, i) are very similar (with deviations of maximum two days, Table S5). This underlines the findings from experiment 2a and 2b, which suggested that moonlight is best perceived in the middle of the night.

### Experiment 4: Modulation of light intensity in simulated natural moonlight impacts the strength of entrainment

Finally, we took into account the modulation of moonlight intensity according to lunar phase, as well as the daily modulation of moonlight intensity. This means that moonlight shifted by 48 min every day, the moonlight intensities gradually changed within a day (moonrise and moonset simulation), and the maximum moonlight intensity changed throughout the lunar cycle (lunar phase simulation). We tested three different moonlight treatments. In the “4-hour low intensity” treatment the total duration of light was reduced to 4 hours, mimicking a narrower gating by the tides and maximum light intensity was down to 0.26 lux (Figure 6b), which corresponds to natural levels of moonlight during full moon. In the “6-hour low intensity” treatment we increased the gating by the tides to six hours with maximum light intensity around 0.24 lux (Figure 6c). Finally, the “6-hour high intensity” treatment had a daily window of six hours of moonlight and the maximum intensity was similar to the standard moonlight treatment (600 lux), i.e. about 2,000-fold brighter than natural moonlight (Figure 6d). We found that all simulated natural moonlight treatments entrain both Vigo-2NM (Figure 6f-h) and Por-1SL (Figure 6j-l) strains, and RI is highly significant for all treatments (Table S5). In the 6h high intensity treatment, Vigo-2NM strain has a very strong artificial peak (Figure 6h), resembling the pattern observed with a six-hour shifting block of high intensity moonlight (Figure 5h). This suggests that despite the intensity modulation, the 6-hour high intensity moonlight is perceived as if it was a 6-hour block of light moving through the night. The picture changes completely in low intensity treatments, where the artificial peak is absent (Figure 6f-h). Interestingly, the 4-hour low intensity treatment, which most closely mimics the natural light regime, shows by far the best entrainment over all experiments (unimodal distribution M2A).

This indicates that low light intensity and light intensity modulation matter for the Vigo-2SL strain, i.e. the Vigo-2SL strain requires and integrates all aspects of this complex moonlight regime. In contrast, the Por-1SL strain is strongly entrained by all natural moonlight treatments (Figure 6j-l). Light intensity does not seem to play a role in strength of entrainment. Notably, the 4-hour low intensity moonlight (Figure 6j) shows stronger entrainment that the 4-hour shifting-block moonlight (Figure 5k). This hints that the modulation of light intensities throughout the lunar months and by the tides may also matter to Por-1SL strain.

As in the shifting-block experiment, we defined day 1 of the 30-day cycle as the day the middle of the moon occurs in the middle of the night, which also matches the day with strongest light intensities (full moon day). Under this definition, for both strains the phase of entrainment matches the standard control moonlight (Figure 6a, e, i), underlining again that both strains seem to be most sensitive to moonlight in the middle of the night.

### Phase-shifts due to timing of light at night might be adaptive and match the natural timing of emergence in Vigo-2NM strain, but not Por-1SL strain

The experiments above (Figures 2-6) show that there is a complex strain-specific integration of several moonlight components that precisely set the phase of entrainment. Interestingly, in the two-hour window in Vigo-2NM strain and four-hour window in Por-1SL strain, we observed phase-shifts depending on the time at night when light was given (Figures 3 and 4). To explore to what extend these phase-shifts are adaptive and if they would lead to emergence during the timing of the low tides in the field, we matched the sensitivity windows to the days in which they are predicted to occur in the field (Figure 7a and b) and compared the emergence patterns to the days of spring tides (Figure 7c and d). Firstly, in both strains the phase-shift of the emergence rhythm compensates for the fact that light at different times of the night occurs on different days in the lunar cycle (Figure 7e and f). In other words, the time of emergence relative to the tidal cycle would stay constant (Figure 7e and f, yellow lines), irrespective of whether light is perceived early in the night and thus early in the tidal cycle, or late in the night and thus later in the tidal cycle (Figure 7e and f, orange lines). This can be considered adaptive, as it makes the phase of emergence robust to weather conditions, which may limit visibility of moonlight to certain times of the lunar cycle in an unpredictable manner.

Secondly, we found that in Vigo-2NM strain emergence under three of the four two-hour windows is predicted to occur a few days before new moon spring tides (compare Figure 7e to 7c), while under one treatment there is an unexpected switch to emergence during full moon (ZT 20-22; see Figure 7e, bottom). The predicted emergence just before new moon corresponds to field observations for new moon populations. Much in contrast, in Por-1SL strain emergence under the four-hour windows does not match the times of spring tides (compare Figure 7f to 7d), contradicting the observed emergence patterns in the field. This may indicate that moonlight entrainment alone is not sufficient for achieving an adaptive phase of emergence in Por-1SL strain.

## Discussion

Moonlight has been shown to be an effective zeitgeber for lunar rhythms for many organisms living in the intertidal region (reviewed in Kaiser and Neumann, 2021), but the effect of the tides on the availability of moonlight in the intertidal zone has rarely been explored. We measured daylight and moonlight in the intertidal zone and found that moonlight is only detectable at wavelengths between 400 and 700 nm (Figure S2a and b). While daylight is detectable at all measured wavelengths (317-953 nm; Figure S3), it is also most intense between 400 and 700 nm (Figure S2c). This is congruent with the fact that moonlight is sunlight reflected in the moon, both having highest intensity at blue wavelengths (400-500 nm) (Cohen et al., 2020). In coastal waters, the highest light intensity of daylight shifts to the green range (500-600 nm) (Mascarenhas and Keck, 2018). The same is true for moonlight, as shown in our measurements (Figure S2a). It remains unclear, if the absence of detectable moonlight below 400 nm and above 700 nm is due to effective filtering of moonlight by the water column, or due to sensitivity limitations of the photometer. Our measurements of light in the intertidal zone also show that highest intensities of both moonlight (Figure 1a) and daylight (Figure S3) are not detected during the zenith of the sun or moon, but during low tide. Additionally, a comparison of our moonlight measurements from the intertidal zone (Figure 1a) to expected moonlight in-land (Figure 1b), clearly show that the duration of detectable moonlight levels is shortened by the tide from >12h to 4-6 hours. This is in line with our experimental observation that short natural moonlight (total of 4 hours) most effectively entrains the Vigo-2SL strain (Figure 6f).

As the timing of the tides varies along the coastline, the occurrence of a low tide in the middle of the night does rarely coincide with the full moon days (it does for Dinard; Figure 1b and c). As a consequence, we can expect from simulations that moonlight availability differs between geographic locations in two major aspects (compare Figures 1c, d, e).

First, light in the middle of the night is present on different days of the moonlight cycle. While in Dinard light in the middle of the night occurs at full moon (Figure 1c), in Vigo it occurs several days after full moon (Figure 1d) and in Port-en-Bessin it occurs several days before full moon (Figure 1e). Indirect evidence that these simulated moonlight patterns match the natural situation come from the observed laboratory emergence phenotypes of *C. marinus* strains from the respective (and other) geographic locations. For each strain, the time between artificial moonlight and emergence in the laboratory corresponds well with the time between days with midnight low tides and spring tide days in the field (Kaiser et al., 2011). That means there are strain-specific phase relationships between moonlight perception and adult emergence which match the local tides. These strain-specific phases are genetically determined and can therefore be considered local adaptations (Kaiser et al., 2011). Additionally, the observation suggests that the relevant moonlight for setting the phase of the rhythm occurs in the middle of the night, congruent with the observations made in our experiments with moonlight windows.

Second, and concerning the very question of the timing of moonlight during the night, our simulations suggest that the highest moonlight intensity occurs at different times of the night for different geographic locations. While in Dinard it occurs in the middle of the night (Figure 1c), in Vigo it occurs a few hours earlier (Figure 1d), and in Port-en-Bessin it occurs several hours later (Figure 1d). Previous experiments with *C. tsushimensis* (Neumann, 1995) and with the Santander strain of *C. marinus* (Neumann, 1969) have shown that there is a window for moonlight sensitivity only during the night. Additionally, the data suggested that this window could have a strain-specific timing. The reason for strain-specific timing of the sensitivity window could be to match the hours of highest moonlight intensity. Alternatively, shifting the timing of moonlight sensitivity could serve to adjust the days during which moonlight is perceived, contributing to adjustment of the phase relationship between light at night and the spring tide days, as described in the paragraph above. In our experiments with different nocturnal moonlight windows (experiments 2a and 2b), the Vigo-2NM strain was most sensitive to moonlight just before the middle of the night (Figure 3h) and the Por-1SL strain was most sensitive in the middle of the night (Figure 4k). So while there are slight differences, the highest moonlight sensitivities (measured as the strongest entrainment of the rhythm) were generally found to occur in the middle of the night and most windows entrained the rhythm. This implies that strain-specificity of the sensitive window is limited. There is no obvious adaptation to the timing of moonlight availability during the night or over the lunar cycle. It underlines that light in the middle of the night is more important than light intensity.

There are two additional lines of evidence in our experiments which suggest that a clear-cut window of moonlight sensitivity is an over-simplistic mechanistic view of moonlight perception, though. First, natural moonlight simulations with naturally low intensity and with modulated light intensities throughout the night and over the lunar cycle resulted in a better entrainment of the rhythm, at least for the lunar rhythm (experiment 4; Figure 6). The observation suggests that the moonlight perception mechanism is complex in that it also uses information coming from the total light intensity (compare Figure 6g to 6h) and from the gradual change of light intensities (compare Figure 6f-g to Figure 5g, h). Second, when moonlight is given in windows throughout the night, both strains responded with changing phase relationships between moonlight perception and emergence peak (Figures 3 and 4). There is not a single sensitivity window coupled to a single fix phase of the emergence peak, but for each strain there is a complex relationship between the timing of light during the night and the phase of the rhythm. Even more, fitting the moonlight windows to the patterns of expected light in the natural habitat (Figure 7) suggests that this flexible phase response to light at night compensates for the fact that the hours of moonlight availability shift through the night over the lunar cycle (see Figure 1). It ensures that the timing of emergence stays constant relative to the lunar cycle, no matter when exactly the moonlight is perceived. This is important in making the timing mechanism robust to weather conditions, which may fully obscure moonlight during parts of the lunar cycle. Such compensation is a remarkable finding, and suggests that the complex response to moonlight is an evolved and fine-tuned property of the moonlight perception mechanism.

Notably, for the Vigo-2SL strain the timing of emergence under the moonlight windows corresponds well with field observations (Figure 7c, e), while for the Por-1SL strain it does not match at all (Figure 7d, f). Thus, for the Por-1SL strain other zeitgebers may be more important than moonlight in adjusting the phase of emergence. This is in line with the observation that the Por-1SL strain does also not seem to be entrained better under the gradual modulation of moonlight (experiment 4; Figure 6). Other zeitgebers known to entrain the semilunar or lunar rhythm of *C. marinus* are water turbulence (Neumann and Heimbach, 1979) and tidal temperature pulses (Neumann and Heimbach, 1984).

The fact that there is a nocturnal window of moonlight sensitivity also gives an elegant solution to the problem of how to discriminate moonlight and sunlight. If the moonlight receptor is only sensitive at night, sunlight will not affect it. Alternatively, or additionally, the discrimination of sunlight and moonlight was suggested to rely on differences in wavelength or light intensities. While we have not tested different wavelengths in our experiments, the fact that the spectral composition of sunlight and moonlight is highly similar (Cohen et al., 2020) argues against a discrimination based on wavelength. We did test light intensities (experiment 1, Figure 2) and found that moonlight is also well perceived at high intensities corresponding to natural daylight, ruling out that the discrimination is based on the fact that moonlight only occurs at low light intensities. Restricting the perception of moonlight to the night remains the most plausible solution in *C. marinus*, although we have shown that the mechanism is more complex than a simple sensitivity window. Notably, in the marine bristleworm *Platynereis dumerilii* a light-receptive cryptochrome (L-cry) adopts different states depending on whether it is excited by bright light (sunlight) or dim light (moonlight). This suggests that in *P. dumerilii* the discrimination of sunlight and moonlight could be based on light intensity (Poehn et al., 2022).

In *C. marinus*, restricting moonlight perception to the night makes entrainment of the lunar rhythm a case of coincidence detection, i.e. the moonlight stimulus is only detected when it coincides with a specific time of day (discussed in detail in Kaiser and Neumann 2021). The fact that moonlight is also perceived in a nocturnal window when it is presented during four days of continuous darkness, suggests that a circadian clock is involved in regulating moonlight sensitivity (Neumann, 1995). We may propose that the circadian clock establishes a complex nocturnal light sensitivity window that ultimately coincides with available moonlight during few nights of the lunar cycle. A similar mechanism of circadian dependence for synchronization of lunar rhythms was recently described for corals, where the spawning rhythm in different species was entrained by different gating windows at night (Komoto et al., 2023).

The molecular mechanism by which moonlight sensitivity is regulated over the day in *C. marinus* remains unknown. Interestingly, the shielding pigment of the larval ocelli appears and disappears over the lunar cycle, making the larval ocelli a “radiometer” without a pigment shield at the time of full moon (Fleissner et al., 2008). The same study also assessed the presence of the shielding pigment at different times of day and did not find a circadian regulation, indicating that regulating the shielding pigment is not the mechanism for restricting light sensitivity to the night.

Our experiments suggest that during the time of moonlight sensitivity, a minimum duration of light at night is required. A moonlight window of 2 hours was not sufficient to entrain the Por-1SL strain (experiment 2a, Figure 3). While the Vigo-2NM strain was entrained by a 2-hour window of light for four nights, it was not properly entrained by a shifting block of moonlight of 2 hours (experiment 3, Figure 5f). This result indicates that not only a certain duration of light during the night is required, but also a certain number of successive nights with the same timing of moonlight availability. It matches the observation that a single night of artificial moonlight is not sufficient to entrain the lunar rhythm (Neumann, 1966). There seems to be a summation or integration of light signals on both the daily and the lunar time-scale.

Finally, our experiments underlined that the second emergence peak observed in the Vigo-2NM strain and other 2NM strains under artificial moonlight (4 nights with continuous bright moonlight) is a laboratory artifact. It is not only absent in the field (Ekrem, Jacobsen, Kokko, Kaiser, in preparation) and under free-run conditions (Neumann, 1966), but also under simulated natural moonlight (Kaiser et al., 2021 supplement and this study). Notably, the natural moonlight simulation presented here is the first laboratory treatment under which both the lunar and the semilunar rhythm of *C. marinus* are precisely entrained without artificial peaks. This lays the basis for QTL mapping for the genetic basis of semilunar (∼15 days) vs. lunar (∼30 days) rhythms, allowing for new avenues of research to finally understand the molecular underpinnings of these enigmatic clocks.

## Acknowledgments

Kerstin Schaefer, Nico Fuhrmann, Alina Waldmann and Miriam Krijewska helped with the experiments. Kristin Tessmar-Raible provided the RAMSES light sensor. Enrique Arboleda retrieved the data from the sensor. The “Biological Clocks” and the “Behavioural Genomics” research groups provided feedback during seminars and discussions. This work was supported by the Max Planck Society via an independent Max Planck Research Group and by an ERC Starting Grant (No 802923) awarded to TSK. CMP was funded by the International Max Planck Research School (IMPRS) for Evolutionary Biology.

## Author Contributions

CMP performed the experiments, analyzed the data and wrote the manuscript. JG and EF deployed and maintained the light sensor in the field. DB conceptualized and performed or supervised the experiments, and edited the manuscript. TSK conceptualized and supervised the experiments, acquired funding, and wrote and edited the manuscript.

## Data availability

Raw light data measured with the RAMSES-ACC-VIS hyperspectral radiometer (TriOS GmbH) placed at Rocher de Bizeux in Dinard, France is available on Max Planck Repository Edmond (https://doi.org/10.17617/3.6OXUES). Processed data other than that provided in supplementary tables was deposited in the same repository.

