## Supplemental_figures for "How light at night sets the circalunar clock in the marine midge *Clunio marinus*"

Supplemental Figure S1. Geographical origin of *C. marinus* strains and locations of light/water level datasets used in this study.

Supplemental Figure S2. Summed wavelengths of moonlight and daylight over four consecutive lunar months measured with radiometer submerged in the intertidal region.

Supplemental Figure S3. Most daylight wavelengths are detected across four consecutive months.

Supplemental Figure S4. Light intensity modulation in the simulated natural moonlight treatments

(a)

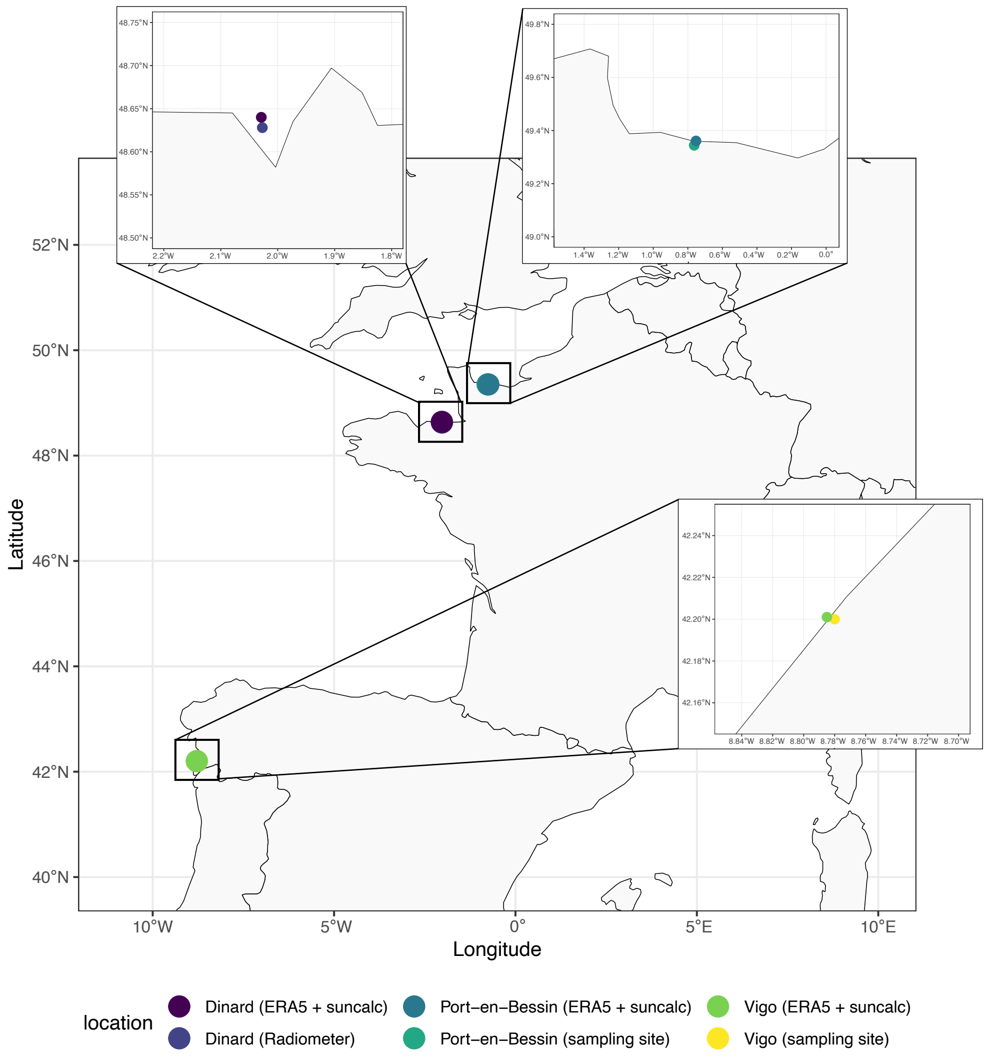


**Supplemental Figure S1. Geographical origin of *C. marinus* strains and locations of light/water level datasets used in this study. (a)** Two populations differing in the period of their lunar rhythm were established and used in this study (Table S1). Vigo-2NM was collected in Vigo, Spain (yellow) and Por-2SL was sampled in Port-en-Bessin, France (dark green). A radiometer was deployed in the intertidal zone of Dinard, France (dark purple) to assess how light intensity and duration is modulated by the tides. Water levels were obtained from the public dataset of ERA5 and relative moonlight data illumination (R package *'suncalc'*) from the same period of time and from close geographical locations (Dinard, light purple, Por, blue and Vigo, light green).

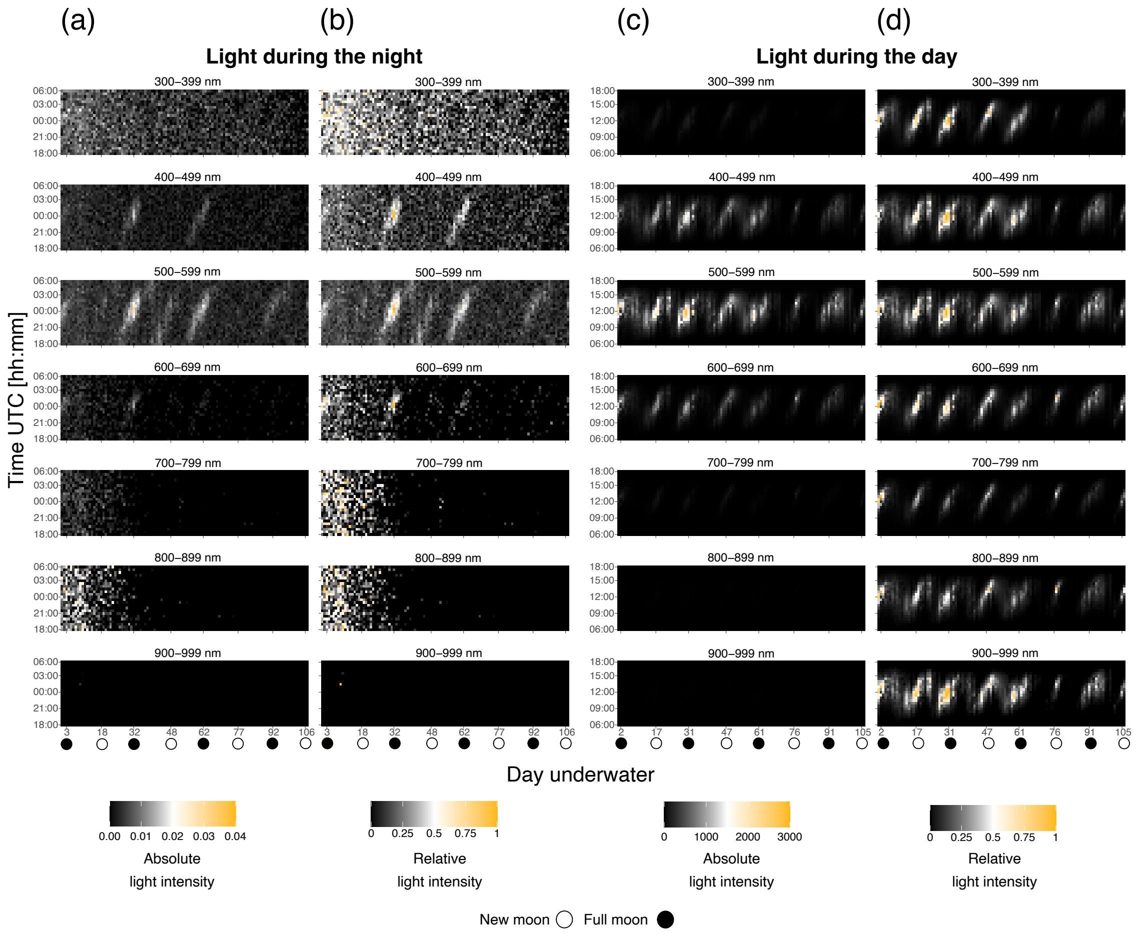


**Supplemental Figure S2. Summed wavelengths of moonlight and daylight over four consecutive lunar months measured with radiometer submerged in the intertidal region.** Heatmaps of light intensity with days shown on the x-axis and time of day on the y axis. Five-minute measurements were averaged every 30 min and 192 wavelengths were summed in 100 nm bins. **(a)** Light detected at night between 18h00 and 6h00. Moonlight can be observed between 400 and 699 nm, with the highest intensity (0.032 mW/m$^{-2}$/nm$^{-1}$) detected in 500-599 nm. 0.04 mW/m$^{-2}$/nm$^{-1}$ was employed to establish the upper threshold for all displayed heatmaps, with the minimum allowed value being fixed at 0. **(b)** Light data from (a) normalized to the highest value detected in each wavelength bin. **(c)** Light detected during the day, between 6h00 and 18h00. Light intensity is strongest in days close to spring tides, with the highest value (2780 mW/m$^{-2}$/nm$^{-1}$) in 500-599 nm. Defining the limits of the heatmaps between 0 and 3000 mW/m$^{-2}$/nm$^{-1}$ shows light detected only between 400 and 699 nm. **(d)** Light data from (c)

normalized to the highest value detected in each wavelength bin. When normalized, it becomes clear that light is detected in all wavelength bins. Differences between light detection in normalized versus non-normalized datasets show that light intensities between wavelengths vary greatly and that moonlight intensity is likely too low for the radiometer to detect wavelengths of lower intensity.

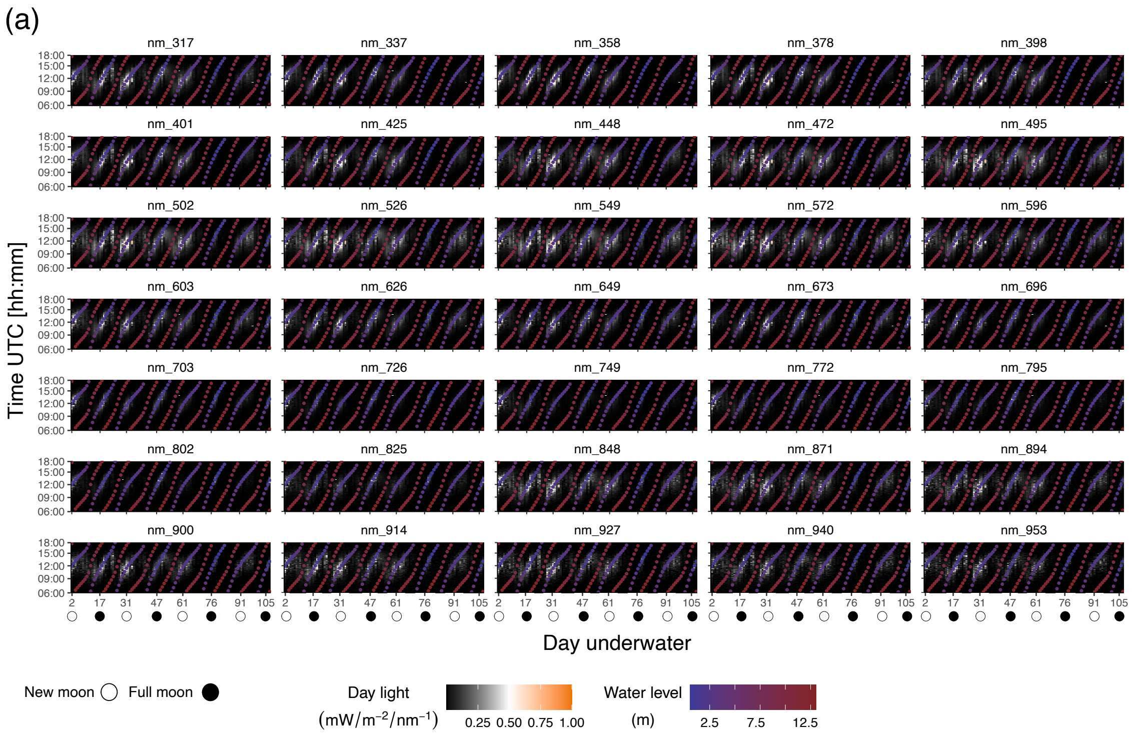


**Supplemental Figure S3. Most daylight wavelengths are detected across four consecutive months. (a)** Heatmaps of five selected wavelengths for each 100 nm bin are shown. Each heatmap is colour-scaled for normalized light intensity (min-max normalization) per wavelength. Days are shown on the x-axis and time of day on the y axis, at a resolution of five-minute measurements. Water levels from the same period of time were obtained from the closest marine station (Saint-Malo) in the publicly available data of maree.info. Highest light intensity at most wavelengths correlates with the timing of low tides.

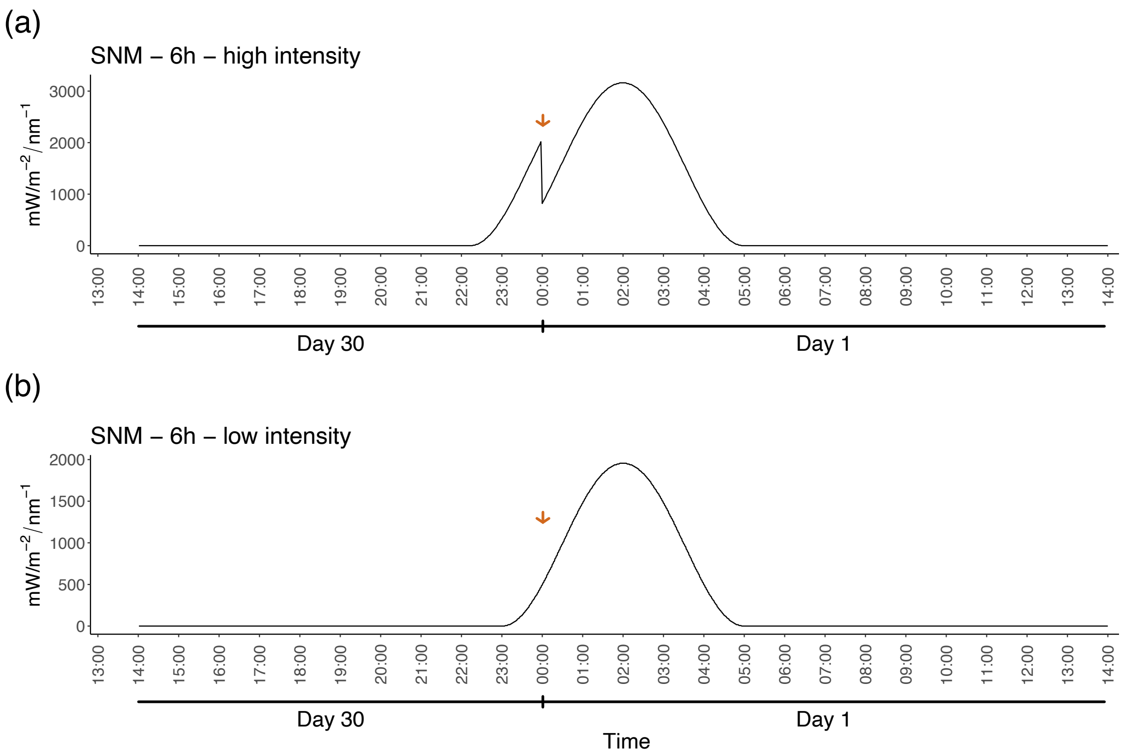


**Supplemental Figure S4. Light intensity modulation in the simulated natural moonlight treatments.** Measurements of light intensity over a full moon night in the simulated natural moonlight programs used in experiment 4. **(a)** "6-hour high intensity" and **(b)** "6-hour low intensity". The Profilux Light computer (GHL, Germany) calculates light intensities per day and fits the 48-minute shift to a 30-day cycle, causing a jump in light intensity at midnight (a, arrow). The recalculation was set to occur during the day in (b) and thus no shift is observed (compare arrows). Note that light measurements values do not correspond to the ones applied in experiment 4 as the Quantum PAR Radiometer was placed close to the LED light with no filters for testing purposes.
